# Whole genome sequencing suggests that “non-pathogenicity on banana (NPB)” is the ancestral state of the *Ralstonia solanacearum* IIB-4 lineage

**DOI:** 10.1101/2022.04.20.488689

**Authors:** Jonathan Beutler, Samuel Holden, Stratton Georgoulis, Darrielle Williams, David J. Norman, Tiffany M. Lowe-Power

**Affiliations:** Department of Plant Pathology, Columbia, Vancouver, University of California Davis, Davis, CA 95616; Land and Food Systems, University of British Columbia, Vancouver, BC, Canada; Department of Plant Pathology, University of California Davis, Davis, CA 95616; UF/IFAS Extension, Mid-Florida Research and Education Center, Apopka, FL 32703

## Abstract

The bacterial wilt pathogens in the *Ralstonia solanacearum* species complex (RSSC) have broad but finite host ranges. Population genetic surveys of RSSC pathogens show that many sequevars (subspecies groups) are predominantly recovered from wilting solanaceous plants. In contrast, strains in the IIB-4 sequevar have been isolated from plants in over a dozen families. Certain IIB-4 lineages have been classified as banana-virulent or “not pathogenic to banana (NPB)”. Prior analysis suggested that the NPB lineage has diverged from the banana-virulent IIB-4 strains. To test this model, we analyzed the phenotypes and phylogeny of a diverse collection of 19 IIB-4 isolates. We used Illumina sequencing to assemble draft genomes of 12 new strains. Based on whole genome phylogenetic analysis, these IIB-4 strains clustered into five subclades. We quantified virulence of each strain on tomato, banana, melon, and impatiens plants. Overall, the virulence patterns correlated with phylogeny. Banana virulence was restricted to the 4/4 IIB-4D subclade (N=4/4 strains) and IIB-4E subclade (N=1/2 strains). Subclades IIB-4D and IIB-4E are sister subclades and their closest relative, the IIB-4A-C subclade, lacked virulence on banana. Our data support a revised model in which banana virulence is an innovation within the IIB4D/E subclades.

**Data Summary:** Illumina sequencing and genome assembly data are available as NCBI BioProject PRJNA826884, and Table S1 lists the accession numbers for assemblies in GenBank and raw sequencing data in SRA. To enable future meta-analyses that identify genetic factors that drive host-range, the raw virulence data is included as Table S2.

## Introduction

The *Ralstonia solanacearum* species complex is a group of related tropical and temperate pathogens that cause plant wilt diseases after they invade and colonize the xylem (Prior et al. 2016; Lowe-Power et al. 2018; Ingel et al. 2022). Collectively, the species complex has a remarkably broad host range, capable of infecting hundreds of host species in dozens of taxonomic families (Hayward 1991; Lowe-Power et al. 2020). However, the breadth in host range is not uniformly distributed across the species complex. Strains in the *R. solanacearum* phylotype IIB sequevar 4 clade (IIB-4) exhibit a particularly broad host range (Ailloud et al. 2015; Wicker et al. 2007) (Lowe-Power et al. 2020). *R. solanacearum* IIB-4 strains have been isolated from 33 plant species in 19 botanical families (Wicker et al. 2007; Norman et al. 2009; Cellier et al. 2012; Hong et al. 2012; Cellier et al. 2015; Gutarra et al. 2017; Ramírez et al. 2020; Zulperi et al. 2014; Lowe-Power et al. 2020; Cellier and Prior 2010). They are distributed throughout Central and South America, the ancestral range of phylotype II *Ralstonia.* This lineage is also globally invasive as IIB-4 strains have become established in Malaysia (Zulperi et al. 2014) and have been imported into Florida (Norman et al. 2009).

*R. solanacearum* IIB-4 have caused several well-studied wilt epidemics. In the 1960s, *R. solanacearum* IIB-4 caused a high-impact Moko disease epidemic of plantains and bananas in Colombia and Peru (French and Sequeira 1970; Buddenhagen and Elsasser 1962). These Moko isolates were also virulent on solanaceous crops: tomato, pepper, and potato (French and Sequeira 1970). In the 2000s, a subgroup of *R. solanacearum* IIB-4 from Martinique were recognized as a novel “not pathogenic to banana (NPB) ecotype” with high virulence on cucurbits but low virulence to banana (Wicker et al. 2007). In 2020, a third ecotype of IIB-4 from Colombia was described (Ramírez et al. 2020); these novel Colombian strains are the only IIB-4 strains that are known to be avirulent on tomato. Overall, the well-documented natural history of IIB-4 strains and the diversity of their hosts position *R. solanacearum* IIB-4 as a model clade for investigating the factors that limit and enable the breadth of host range in bacterial wilt pathogens.

Our prior comparative genomic analysis of 14 phylotype II strains, including 5 IIB-4 strains, was consistent with a hypothesis that the NPB lineage diverged from the banana-virulent IIB-4 strains. To test this hypothesis, we collected IIB-4 isolates from additional sources, inferred their phylogeny by whole-genome sequencing and assembly, and analyzed their virulence phenotypes on host species from four botanical families: banana, tomato, melon, and impatiens.

## Results

We quantified the natural variation in virulence of 19 *Ralstonia solanacearum* IIB-4 strains against four natural host species from four botanical families. The strains were isolated from diverse plants and locations (see Table S1). Seven strains were isolated in Peru and Colombia from 1960-1963. Of these, six were from plantain during an epidemic of Moko disease, and one from *Heliconia* (Buddenhagen and Elsasser 1962; French and Sequeira 1970). Three strains were isolated on *Anthurium* and *Heliconia* during the Martinique epidemic (Wicker et al. 2007). Seven strains were isolated from ornamentals imported into Florida. Of these, six were isolated between 1996-2012 from *Pothos,* and one from *Anthurium* in 1997 (Norman et al. 2009). One strain was isolated in 1999 from cucumber in Brazil (Ailloud et al. 2015), and one in 2017 from *Pothos* in the Dominican Republic.

To determine the diversity of these strains, we used Illumina sequencing to assemble draft genomes of 12 strains. An additional four unpublished genomes were shared by Caitilyn Allen and Boris Vinatzer. The draft genome assembly of P488 is lower quality than others. The P488 assembly is fragmented into 200 contigs with an N_50_ of ~56 kb while the other assemblies have 40-115 contigs with an N_50_ of 111-414 kb. Full genome assembly statistics are in Table S1. We collected genomes of five additional IIB-4 strains, the closely related IIB-51 strain CFBP7014, and two distantly related strains (phylotype I GMI1000 and phylotype IIC K60 (Sharma et al. 2022)) from NCBI.

We constructed a phylogenetic tree using 4,317 genes comparing 369,907 intragenic SNPs (Fig 1). Three of the IIB-4 genomes had longer branch lengths than expected (P488, CFBP6783, and UW179), consistent with the lower-quality, more fragmented assemblies for these strains. The tree separated the IIB-4 strains into five branches, which we refer to as subclades A to E. The *Heliconia* isolate from 1961 (UW170) was the sole genome on subclade A. Seven *Pothos* and *Anthurium* isolates from Florida, and one from the Dominican Republic all clustered in subclade B. The Brazilian cucumber isolate IBSBF1503 clustered into subclade C with the three strains from the Martinique epidemic. The four Peruvian strains from the 1960s Moko epidemic clustered into subclade D. Subclade E contained three Colombian strains isolated from plantain in the 1960s, three Colombian strains recently isolated from plantain, and one Mexican strain isolated from potato.

**Fig 1.**
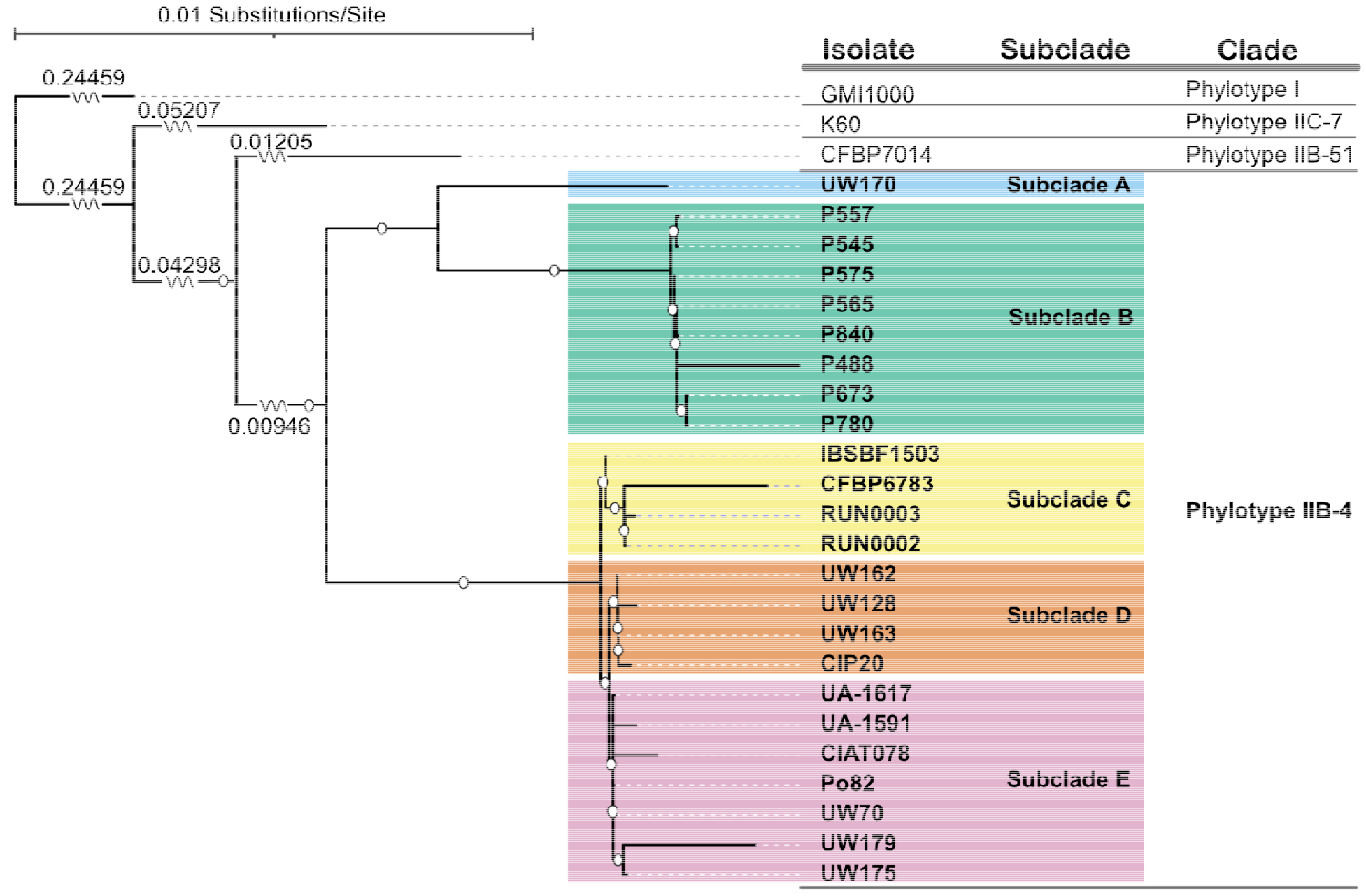
Maximum likelihood tree describing 27 strains of the *Ralstonia solanacearum* species complex, based on 369,907 intragenic SNPs. The tree includes 24 IIB-4 strains and three reference strains from phylotype IIB-51 (CFBP7014), IIC-7 (K60) (Sharma et al. 2022), and I (GMI1000). Strains are described as belonging to one of five IIB-4 subclades (A-E). 95%bootstrap support is indicated by a white circle. Some branch lengths were condensed for clarity. SNPs were identified by grouping orthologous genes across all isolates using PIRATE v1.0.4 with default parameters and nucleotide-based clustering (Bayliss et al. 2019). Orthologous gene groups identified by PIRATE were concatenated and reduced to positions present in at least 20 accessions, with at least one variable nucleotide, by a python script Phylip_reducer.py (github.com/Erysiphe-graminis/Utilities). These positions were used to generate the tree using RAxML v8.2.12 with the GTRGAMMA model, Lewis ascertainment bias correction, and 1000 bootstraps (Stamatakis 2014). GMI1000 was the outgroup. The tree was visualized using ITOL v6.5.2 (Letunic and Bork 2021), and the figure prepared using Adobe Illustrator. To evaluate the impact of excluding infrequently occurring genes, two additional trees were generated using the same parameters except that SNP positions could be present in (1) at least 10 accessions, or (2) in any number of accessions. These trees were topologically identical with subtle variations in lengths of some branches.

We quantified virulence of 19 IIB-4 strains on four diverse plant species that are documented natural hosts for *Ralstonia* IIB-4: tomato (*Solanum lycopersicum* cv. Moneymaker), melon (*Cucumis melo* cv. Sweet Granite), impatiens (*Impatiens walleriana* cv. Beacon Orange), and banana (AAA *Musa acuminata* cv. Dwarf Cavendish). Tomato, melon, and impatiens were grown from seed while banana plants were purchased as tissue cultured plantlets. Plants were grown in growth chambers at 28°C (tomato, melon, and impatiens) or 32°C (banana) with a 12 hr light cycle and 40% humidity. Tomato and melon seeds were sown into Sunshine Mix #1 to a depth of 1 cm, and impatiens were sown onto the surface of Sunshine Mix and topped with 1 mm vermiculite. Seedlings were transplanted into 3.5” pots with approximately 60 g Sunshine Mix at 7 days (melon), 14 days (tomato), 21 days post sowing (impatiens), or at the 3-leaf stage (banana).

We inoculated plants at 21 days post sowing (tomato and melon), 35 days post sowing (impatiens), or at the 4-leaf stage (banana). We inoculated tomato and impatiens via a standard cut-petiole approach where the lowest petiole was excised, and a 2 μl droplet of bacterial suspension (containing approximately 1000 CFU) was placed on the cut surface (Khokhani et al. 2018). Due to rapid wound responses of cucurbits (Zimmermann et al. 2013), we inoculated melons by applying a 2 μl droplet of 1000 CFU onto a razor blade and excising a petiole with the infested blade. Banana plants were inoculated by drenching the soil with a bacterial suspension (50 mL with 5×10^7^ to 1×10^8^ CFU/g soil) after the roots were lightly wounded with a 2 cm-wide metal spatula (Ailloud et al. 2016). We rated symptoms daily for 14 days (Fig. 2). We rated tomato symptoms on a standard 0-4 disease index (DI) where 0 = no wilt, 1 = 0.1-25% of leaflets wilted, 2 = 25.1-50% of leaflets wilted, 3 = 50.1-75% of leaflets wilted, and 4 = 75.1-100% of leaflets wilted (Khokhani et al. 2018). We measured symptoms on melon, impatiens, and banana as the percentage of leaves with wilt. Symptom measurements, area under the disease progress curve (AUDPC), and incidence are displayed in Table S2.

**Fig 2.**
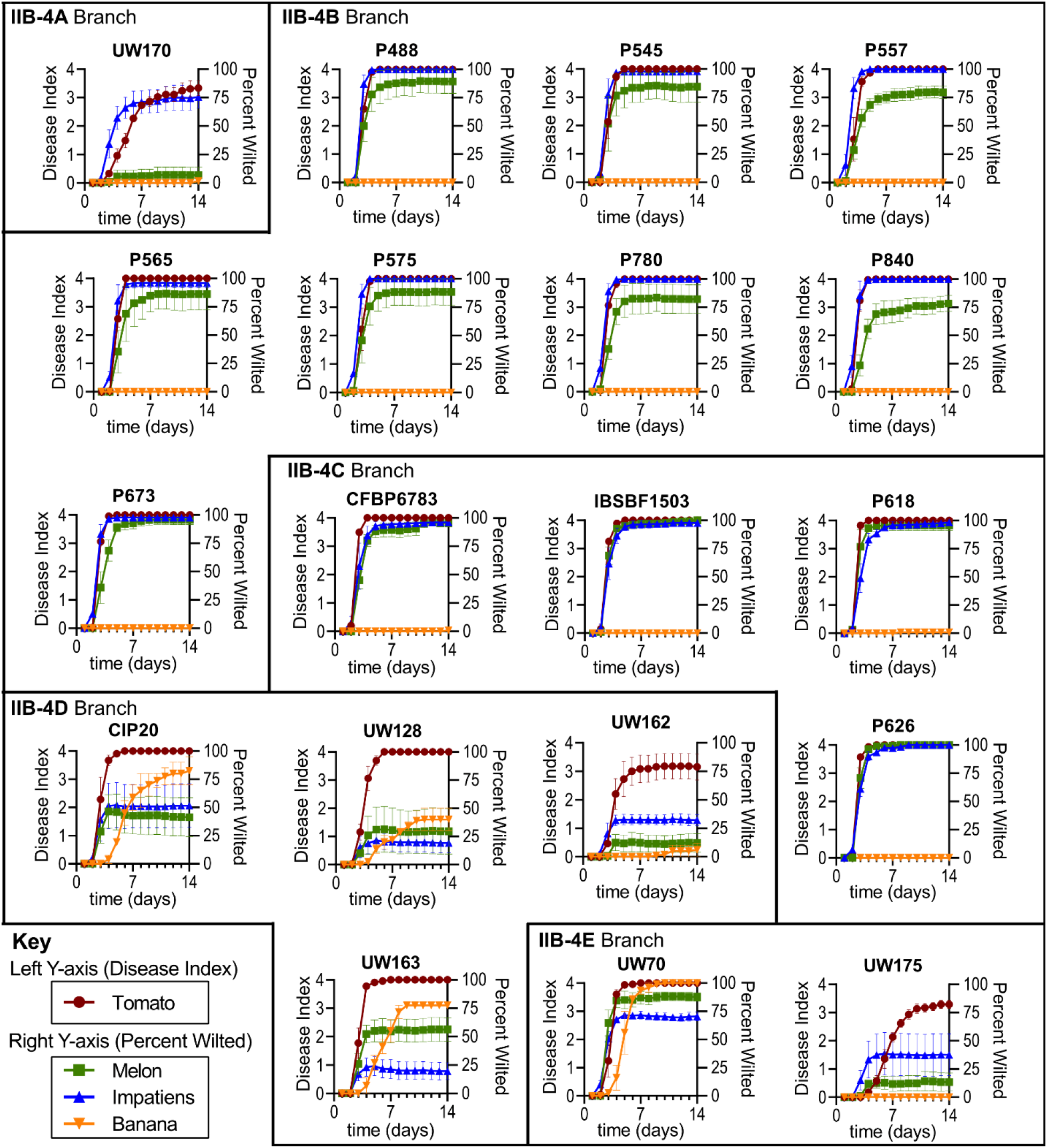
Disease progress of 19 *R. solanacearum* IIB-4 strains on four host species: tomato cv. Moneymaker (n≥44 plants), melon cv. Sweet Granite (n≥43), impatiens cv. Beacon Orange (n≥44), and Banana cv. Dwarf Cavendish (n≥19). Each graph represents one strain. Virulence on tomato was quantified on 0-4 disease index (left Y-axes), and virulence on other hosts was quantified as a percentage of leaves with wilt (right Y-axes). Symbols represent the mean of 2-3 trials and bars show standard error of the mean (SEM). Strains are grouped based on the phylogeny from Fig 1. Raw virulence data is available in Table S2.

Although all strains were pathogenic to tomato, three strains had reduced virulence: UW170 (subclade A), UW162 (subclade D), and UW175 (subclade E). Because these strains have been stored as “waterstocks” (in distilled water at room temperature) for decades, we suspect that these strains acquired mutations in culture storage that reduced their virulence. The subclade A strain UW170 was moderately virulent on both tomato and impatiens but caused minor wilt on only a few banana or melon plants. The subclade D strain UW162 had moderate virulence on tomato and mild virulence on impatiens, melon, and banana. The subclade E strain UW175 had moderate virulence on tomato, mild virulence on impatiens and melon, and no virulence on banana.

Virulence of strains generally correlated with their phylogenetic relationships. The subclade B and C strains lacked virulence on banana but highly virulent on tomato, impatiens, and melon. The subclade C strains displayed slightly higher virulence on melon than the subclade B strains. Three other subclade D strains were highly virulent on tomato and had low-to-moderate virulence on impatiens, melon, and banana. The subclade E strains included UW70 which was highly virulent on tomato, melon, and banana. UW70 was more virulent on impatiens than subclade D strains, but less virulent than the subclade B and subclade C strains.

## Discussion

Pathogen populations are collections of genotypically and phenotypically diverse individuals. By inferring phylogenetic relationships of strains, we can set the groundwork for genome-wide analyses that associate genotypic differences with phenotypes. For plant pathogenic *Ralstonia,* the sequevar system has been widely used to classify strains to sub-species groups (Fegan and Prior 2005; Lowe-Power et al. 2020). Although the sequevar system is merely based on sequence alignments of a single 750 bp DNA marker, we recently validated that sequevars for *Ralstonia* phylotype II (i.e. *Ralstonia solanacearum*) accurately reflect the strains’ core genome phylogenic trees (Sharma et al. 2022). However, the sequevar system broke down for the phylotype I branch of *Ralstonia pseudosolanacearum* (Sharma et al. 2022), which is known to be highly recombinogenic (Wicker et al. 2012; Peeters et al. 2013). Moving forward, bacterial taxonomy will continue to be influenced by whole-genome comparisons (Parks et al. 2020). Taxonomic systems like the Life Identification Number (LIN) system (Tian et al. 2020; Vinatzer et al. 2017) are likely to supersede the sequevar system. LINs are a systematic taxonomy based on multiple levels of whole genome average nucleotide identity (ANI). Strains in the IIB-4 sequevar have pairwise ANI of over 99.5% on their shared genes, and they are circumscribed in the LIN group 14_A_1_B_0_C_0_D_0_E_3_F_0_G_0_H_1_I_0_J_0_K_0_L_0_M_ (Sharma et al. 2022).

Although many *Ralstonia* isolates only wilt plants efficiently at tropical temperatures, some *Ralstonia* isolates can wilt plants at temperate temperatures. Cool-virulence is famously a trait of the global pandemic lineage of *Ralstonia* (IIB-1 AKA the U.S. Select Agent-regulated “Race 3 Biovar 2” group)(Champoiseau et al. 2009), but recent studies have shown that multiple lineages *of Ralstonia* can wilt tomato plants at cool temperatures: IIB-27, III-48, and the focal clade of this study, IIB-4 (Bocsanczy et al. 2014, 2017). Although we carried out this study’s virulence assays at warm, tropical temperatures, several of our isolates, P673 (subclade B), CFBP6783 (subclade C), and UW163 (subclade D) can wilt tomatoes at cool temperatures (18°C) (Bocsanczy et al. 2014, 2017).

Early studies suggested that IIB-4 strains have binary host range phenotype with either a “Moko” phenotype with virulence on banana/ tomato/ plantain or a “NPB” phenotype with virulence on tomato/ plantain/ anthurium/ cucurbits (Ailloud et al. 2015; Wicker et al. 2007; Ailloud et al. 2016). However, our study shows that the virulence patterns are more complex. First, we show that banana virulence is rare among the strains tested. The NPB phenotype was shared by the subclade C strains previously described as “not pathogenic to banana” (Ailloud et al. 2015) and strains from subclade A and B. Our results suggest that virulence on banana might be an acquired trait within the subclade D/E lineage. Consistently, virulence on banana is a polyphyletic trait shared by distantly related lineages in phylotypes IIA, IIB, and IV (Ray et al. 2021; Albuquerque et al. 2014; Obrador-Sánchez et al. 2017). Our study did not include any genomes from the recently identified IIB-4 strains from Colombia that lacks virulence on tomato (Ramírez et al. 2020). If those strains branch independently from the five subclades of the 24 IIB-4 strains in this study, we propose to name the newly described strains as “subclade F”.

One subclade E strain (UW70 also known as “CIP417”) was highly virulent on all host species tested. UW70 was isolated from plantain in Colombia in the 1960s and is closely related to strains isolated from plantain in Colombia in 2012 (UA-1591 and UA-1617) (Ramírez et al. 2020). Interestingly, UW70 was highly virulent on melons, but UA-1591 is reported to be avirulent on cucumber (Ramírez et al. 2020). Cucurbit virulence has also been documented in phylotype I strains isolated in Taiwan, Thailand, China, and Japan (He et al. 2021; Yahiaoui et al. 2017; Horita et al. 2014; Lin et al. 2014).

Traits of several of this study’s isolates were studied: subclade C strain CFBP6783, subclade D strain UW163 and subclade E strain UW70. Lebeau et al. (2011) measured virulence of CFBP6783 and 11 other diverse *Ralstonia* strains against 30 accessions of tomato, eggplant, or pepper accessions in the “Core-TEP” panel. In this study, CFBP6783 exhibited a unique pattern of high virulence on 10/10 tomato, 10/10 pepper, and 6/10 eggplant accessions. CFBP6783 is virulent on plantain (AAB *Musa* sp. cv. Dominico-Harton)(Wicker et al. 2007) and potato (cv. Bintje Munterschen × Franschen)(Cellier and Prior 2010), but is not virulent on Cavendish banana (Wicker et al. 2007 and this study). UW70 and UW163 were highly virulent on *Musa balbisiana* Colla, tomato (*Solanum lycopersicum* L. ‘Bonny Best’), pepper (*Capsicum annuum* L. ‘Sweet Yellow’), and potato (*Solanum tuberosum* L. ‘Russet Burbank’)(French and Sequeira 1970). In contrast, they exhibited low virulence on eggplant (*Solanum melongena* L. ‘Black Beauty’) and tobacco (*Nicotiana tobacum* L. ‘Bottom Special’). In a separate study, UW163 (D subclade) and UW179 (E subclade) lacked virulence on Anthurium while subclade C strains CFBP6783 and IBSBF1503 were virulent on Anthurium (Ailloud et al. 2015).

Our findings reinforce the long-recognized phenotypic plasticity within the *Ralstonia solanacearum* species complex (Buddenhagen 1985). In addition to host range variation within sequevar IIB-4, banana-virulent strains in IIA-24 and IIA-41 clades vary in their virulence on tomato (Albuquerque et al. 2014). A study of phylotype I *Ralstonia* from Guanxi, China showed that closely related strains shared virulence on tomato, eggplant, tobacco, and potato but varied in their virulence on pepper, mulberry, ginger, and cucurbits (He et al. 2021).

In attempts to classify strains by their host range, scientists have attempted to sort *Ralstonia* into “Races”, “Pathoprofiles”, “Pathotypes”, and “Ecotypes” (Lebeau et al. 2011; Cellier et al. 2012; Buddenhagen 1985; Peeters et al. 2013). Having clear phenotypic classifications would be both intrinsically satisfying and immensely helpful for clarifying regulations. In the future, this may be possible through sophisticated statistical analysis. However, these data science approaches will require a concerted and sustained effort to generate robust data on the genomic and phenotypic diversity of the global *Ralstonia* population. In parallel, a systematic database should be developed to consolidate and communicate knowledge on host range of different *Ralstonia* lineages. Continued study is needed to gain deeper insight into the genetic and evolutionary drivers of host range in the *Ralstonia solanacearum* species complex

## Acknowledgements

We thank Caitilyn Allen for sharing strains used in this study and Caitilyn Allen, Boris Vinatzer, and Parul Sharma for sharing unpublished genomes. Work was funded by a Henry A. Jastro Graduate Research Award to J. Beutler; the UC Davis College of Agricultural and Environmental Sciences and USDA CA-D-PPA-2610-H to T. Lowe-Power; the Plant Agricultural Biology Graduate Admissions Pathways Program to D. Williams; the USDA Floriculture and Nursery Research Initiative and the University of Florida Institute of Food and Agricultural Sciences to D. Norman.

